# Low affinity membrane transporters can increase net substrate uptake rate by reducing efflux

**DOI:** 10.1101/213140

**Authors:** Evert Bosdriesz, Meike T Wortel, Jurgen R Haanstra, Marijke J Wagner, Pilar de la Torre Cortés, Bas Teusink

## Abstract

Cells require membrane-located transporter proteins to import nutrients from the environment. Many organisms have several similar transporters for the same nutrient, which differ in their affinity. Typically, high affinity transporters are expressed when substrate is scarce and low affinity ones when substrate is more abundant. The benefit of using low affinity transporters when high affinity ones are available has so far remained unclear. Here, we investigate two hypotheses. First, it was previously hypothesized that a trade-off between the affinity and the maximal catalytic rate explains this phenomenon. We find some theoretical and experimental support for this hypothesis, but no conclusive evidence. Secondly, we propose a new hypothesis: for uptake by facilitated diffusion, at saturating extracellular substrate concentrations, lowering the affinity enhances the net uptake rate by reducing the substrate efflux rate. As a consequence, there exists an optimal, external substrate concentration dependent transporter affinity. An *in silico* analysis of glycolysis in *Saccharomyces cerevisiae* shows that using the low affinity HXT3 transporter instead of the high affinity HXT6 enhances the steady-state flux by 36%. We tried to test this hypothesis using yeast strains expressing a single glucose transporter that was modified to have either a high or a low affinity. Due to the intimate and reciprocal link between glucose perception and metabolism, direct experimental proof for this hypothesis remained inconclusive in our hands. Still, our theoretical results provide a novel reason for the presence of low affinity transport systems which might have more general implications for enzyme catalyzed conversions.

## 1 Introduction

Cells need to acquire all their nutrients and energy sources from the environment. Since hardly any of these can diffuse freely through the membrane, nutrient uptake requires transporter proteins. Often, a cell has several different transporters for the same nutrient. A recurring phenomemon is that these different transporters have different affinities. For example, the yeast *Saccharomyces cerevisiae* has 17 different glucose transporters [1], with affinities ranging from *K*_*M*_ ≈ 1 mM for the highest to *K*_*M*_ ≈ 100 mM for the lowest affinity transporters. Other examples of nutrient transport by both high and low affinity transporters are glucose uptake in human cells transporter [2] and in *Lactococcus lactis* [3], phosphate and zinc uptake in yeast [4], and lactate transport in mammalian cells [5, 6]. The *Arabidopsis* nitrate transporter CHL1 was shown to be able to switch between a high and low affinity mode of action through phosphorylation of the protein [7]. Typically, the high affinity transporters are expressed under conditions of low substrate availability and the low affinity transporters when substrate is plentiful. While the benefit of employing a high affinity transporter under substrate scarcity is evident, the reason for switching to low affinity transporters when substrate is more abundant remains unclear. Why is a high affinity transporter not always preferable over a low affinity one?

Previously, several hypotheses have been suggested to explain the benefit of using low affinity carriers. One hypothesis states that these increase the ability of cells to sense extracellular substrates. Levy et al. convincingly show that low affinity carriers for phosphate and zinc allow the cell to sense depletion of phosphate and zinc early, and consequently, the cells can adapt their physiology to a phosphate or zinc-poor environment [4]. However, for substrates with a higher import rate, such as glucose, there might be a stronger selection pressure on efficiency of uptake than on accurate sensing. Moreover, or perhaps consequently, separate extracellular substrate sensors have been described (for example in *S. cerevisiae* [1]), and glucose sensing and uptake in *S. cerevisiae* are known to be uncoupled [8].

A second hypothesis is related to transporter efficiency, and states that there is a trade-off between the affinity and specific activity (*k*_*cat*_) of a transporter (suggested by Gudelj et al.[9] based on data by Elbing et al. [10]). The cell membrane is a crowded place, and valuable space taken up by a particular membrane protein cannot be used by another one [11]. This implies that, besides the expenditure of energy and precursor metabolites, expression of membrane transporter proteins entails an additional cost, and there is a strong selective pressure to maximize the uptake flux per unit transporter expressed in the membrane. Under high substrate availability, it is beneficial to switch to low-affinity, high-*k*_*cat*_ transporters. While there is some theoretical support for a rate-affinity trade-off for particular reaction schemes [12, 13], this depends on untestable assumptions about the free energy profile and it does not apply to typical reaction schemes of membrane transport processes, such as facilitated diffusion. We will study the theoretical and experimental basis of this trade-off for transport by means of facilitated diffusion.

These hypotheses do not convincingly explain the presence of many distinct carriers in, for example, *S. cerevisiae*. In this paper, we present a new hypothesis based on the reversibility of the transport process. We focus in particular on facilitated diffusion, an often occurring mechanism of which glucose uptake in yeast and human cells are examples. We suggest that an increased affinity not only increases the rate of the substrate influx into the cell, but also that of substrate efflux out of the cell. Due to the nature of a substrate carrier, the impact of the outward transport remains significant even at saturating extracellular concentrations [14]. Therefore, we cannot neglect the outward transport solely because extracellular concentrations are much higher than intracellular concentrations. We argue that the main difference between high- and low-affinity facilitated-diffusion transporters is that for the high-affinity transporters often both sides are saturated with substrate, while for the low-affinity transporter only the extracellular side is. At the same (high) extracellular substrate concentration the low-affinity transporter would therefore have a higher import activity. This reasoning is graphically depicted in Figure 1. With this hypothesis we challenge the intuitive assumption that a higher affinity always leads to a higher net uptake rate.

**Figure 1:**
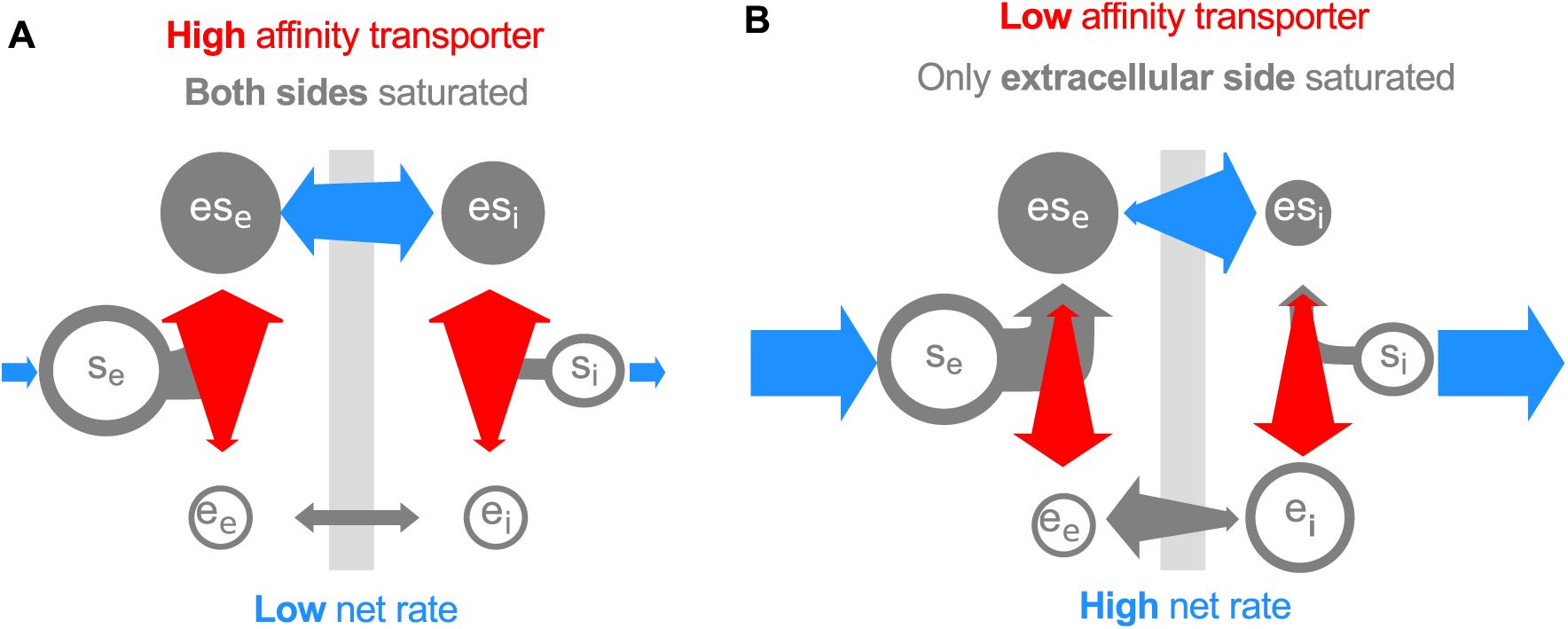
Lower affinity can enhance uptake by reducing substrate efflux. Both panels depict a situation with a high extracellular and moderate intracellular substrate concentration. **A** A high affinity transporter will cause both the inward facing and the outward facing binding sites to be predominantly occupied, i.e., the transporter is saturated with substrate on both sides of the membrane (both *es*_*e*_ ≫ *e*_*e*_ and *es*_*i*_ ≫ *e*_*i*_). As a result, the efflux rate will be nearly as high as the influx rate, and the net uptake rate is very low. **B** Reducing the affinity of the transporter reduces the saturation of the transporter at the intracellular side. Provided *es*_*e*_ is high enough, the transporter will still be saturated at the extracellular side. The efflux will be reduced and hence the net uptake rate increases.

## 2 Results

### 2.1 Mathematical model of facilitated diffusion kinetics

Transport of substrate *s* over a membrane by means of facilitated diffusion can generally be described by a four-step process (depicted in Figure 2). These steps are: (i) extracellular substrate *s*_*e*_ to carrier binding, (ii) transport of *s* over the membrane, (iii) release of *s* in the cytosol and (iv) return of the substrate-binding site to the periplasm-facing position. Note that step (iv) is the only step that distinguishes this scheme from reversible Michaelis-Menten kinetics. We will later discuss the significance of this distinction.

**Figure 2:**
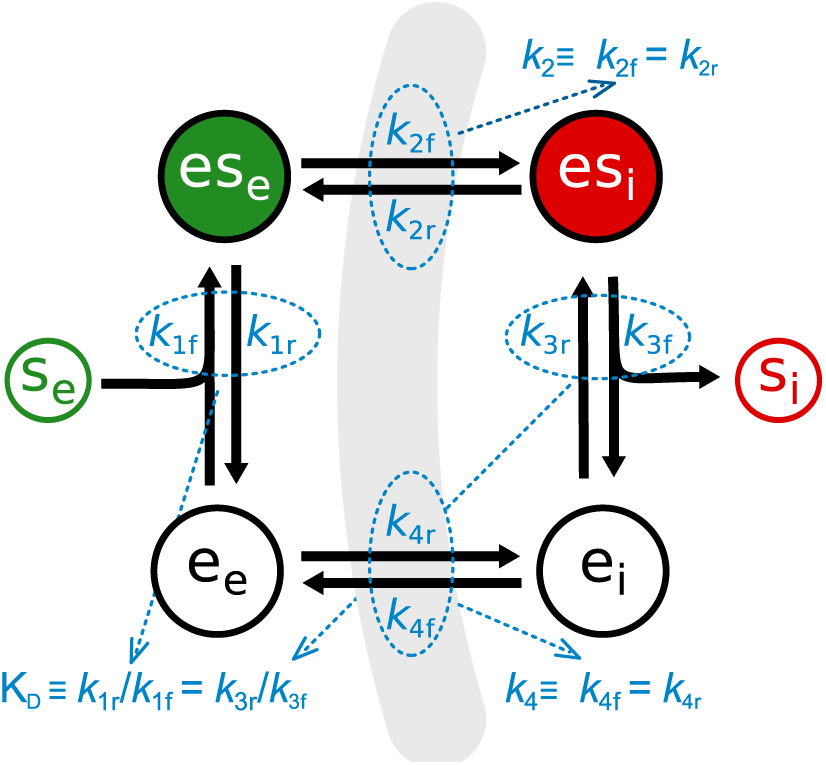
Model of nutrient uptake by facilitated diffusion. In this model the transporter *e* switches between conformations with an inward-facing (*e*_*i*_) and outward-facing (*e*_*e*_) substrate binding site. When this conformation change takes places with an occupied substrate binding site (*es*_*e*_ or *es*_*i*_), this results in translocation of the substrate over the membrane. All steps are reversible and the state transitions rates are given by mass action kinetics. Throughout the main text, we make the assumption that binding is much faster than transport and that the carrier is symmetric. The former assumption allows us to use the quasi steady-state assumption for substrate to transporter binding, *i.e. e*_*x*_ and *es*_*x*_ are in equilibrium, with dissociation constant *k*_*D*_ ≡ *k*_1*r*_ /*k*_1 *f*_ = *k*_3 *f*_ /*k*_3*r*_. The latter assumption implies *k*_2 *f*_ = *k*_2*r*_ ≡ *k*_2_ and *k*_4*f*_ = *k*_4*r*_ ≡ *k*_4_

Thermodynamics dictate that all individual steps are reversible. Moreover, there is no energy input in this transport cycle, so the equilibrium constant *K*_*eq*_ = 1. For convenience, we will make two biologically-motivated assumptions that considerably simplify the rate equation in terms of the first order rate constants. However, relaxing these assumptions does not qualitatively alter our conclusions (cf. Supplementary Information). The assumptions are: (a) binding and unbinding of the substrate to the transporter is much faster than transport of the substrate over the membrane, i.e., binding is assumed to be in quasi steady state, and (b) the transport process is symmetrical. This implies two things, the intraand extracellular substrate-transporterdissociation constants are equal and the forward and reverse rate constants of steps (ii) and (iv) are equal, i.e., *k*_2 *f*_ = *k*_2*r*_ ≡ *k*_2_ and *k*_4 *f*_ = *k*_4*r*_ ≡ *k*_4_. The latter assumption is sensible because the “substrate” and the “product” of the transportation step are chemically identical. Therefore, there is no a priori reason why e.g., extracellularly the substrate should bind tighter to the carrier than intracellularly. The consequence of this assumption is that the *K*_*M*_ of substrate influx and substrate efflux are equal. For e.g., hexose transport in *S. cerevisiae* this is indeed the case for all 7 Hxt carriers that were characterized by Maier et al. [15] (also see [10]). The rate equation takes the form

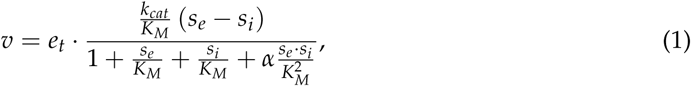

with

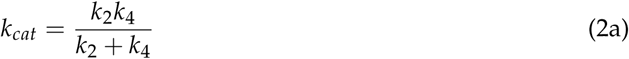

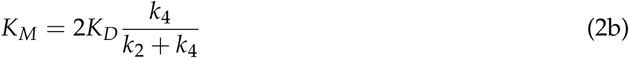

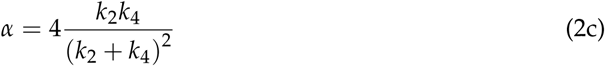

Where 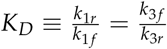 is the dissociation constant of transporter-substrate binding.

### 2.2 The trade-off hypothesis: The theoretical basis of the trade-off between rate and affinity

A comparison of equations (2a) and (2b) immediately shows both the strength and the weakness of the hypothesis that there is a trade-off between the *k*_*cat*_ and the affinity of a transporter. Since both 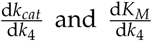 are always positive, any mutation that increases *k*_4_ enhances the *k*_*cat*_ but reduces the affinity. In that sense, there might be a rate-affinity trade-off. However, any mutation that decreases the *K*_*D*_ enhances the transporters affinity without affecting the *k*_*cat*_, and since 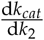 is always positive but 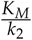 is always negative, an increase in *k*_2_ enhances both the affinity and the *k*_*cat*_. This means that only if *k*_2_ and *K*_*D*_ are constrained by biophysical limitations, there is a real trade-off between rate and affinity. Whether or not this is the case is generally difficult to establish. An additional confounding factor is the α-term, which describes the asymmetry between an occupied and an unoccupied carrier and which is affected by any mutations affecting *k*_2_ and *k*_4_. While α does not affect the *k*_*cat*_ and *K*_*M*_, it does affect the uptake rate. The higher it is (i.e., the larger the asymmetry), the lower the uptake rate.

When the assumptions of fast substrate to transporter binding and symmetry are dropped, an analytic evaluation of this trade-off becomes infeasible. The macroscopic kinetic parameters depend in a complicated way on the first order rate constants, and the latter are interdependent. A parameter sampling approach indicates that also in this case there are some rate-constants for which a trade-off can be found, while for others this is not the case (Figure S1). We refer to the Supplementary Information for a more detailed discussion.

### 2.3 The reduced efflux hypothesis: A lowered affinity can enhance the net uptake rate by reducing substrate efflux

When discussing the transporter’s *k*_*cat*_ and affinity in terms of a trade-off, one makes the implicit assumption that for flux maximization a high affinity is, all else being equal, always better than a low affinity. While this might sound like an obvious statement it is, in fact, not true; decreasing the affinity of the transporter without affecting the *k*_*cat*_ can actually enhance the net uptake rate.

To see why, consider a high affinity transporter fully saturated with extracellular substrate. Now, suppose that the intracellular substrate concentration is much lower than the extracellular, but still well above the *K*_*M*_ of the transporter, i.e. *s*_*e*_ ≫ *s*_*i*_ ≫ *K*_*M*_. In this situation, each time the transporter moves its binding site over the membrane a substrate molecule will be transported, regardless of whether it moves from the outside to the inside, or the other way around. Hence, despite a considerable concentration gradient, the influx and efflux rates will be nearly equal, and the net uptake rate will be close to zero. In other words, the transporter is severely inhibited by its product. In contrast, consider the same situation, but now with a low-affinity transporter, which has a *K*_*M*_ above the intracellular glucose concentration, but still well below that of extracellular glucose, *s*_*e*_ ≫ *K*_*M*_ > *s*_*i*_. In this case, the transporter will operate close to its *V*_*max*_, because it’s forward rate is saturated, but the efflux rate is low. This reasoning is graphically depicted in Figure 1.

The argument above implies that there is a condition-dependent optimal affinity, 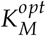 for the transporter, which maximizes the net uptake rate as a function of intracellular and extracellular substrate concentration. This is given by (cf. Supplementary Information):

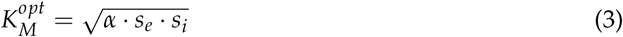

Figure 3A (solids lines) visualize this relation between the *K*_*M*_, the steady-state uptake rate *J*, and the external substrate concentration *s*_*e*_. Below a certain value, reducing the *K*_*M*_ reduces *J*, and as *K*_*M*_ approaches 0, so does *J*. At which *K*_*M*_-value the maximal *J* is attained depends on *s*_*e*_. This is in stark contrast to reversible Michaelis-Menten kinetics, where an increased affinity always increases the uptake rate (Figure 3A, dashed lines). As noted previously, the fundamental difference between these two models is the step between intracellular substrate release and relocation of the binding site to the extracellular side of the membrane. In the carrier model, intra- and extracellular substrates are not directly competing for the same binding sites, and the substrate efflux rate is in effect insensitive to the extracellular substrate concentration. This explains the qualitatively different behavior of the two kinetic schemes. The result that there is an optimal affinity of the transporter also holds for the non-symmetric carrier models (cf. Supplementary Information and Figure S2)

**Figure 3:**
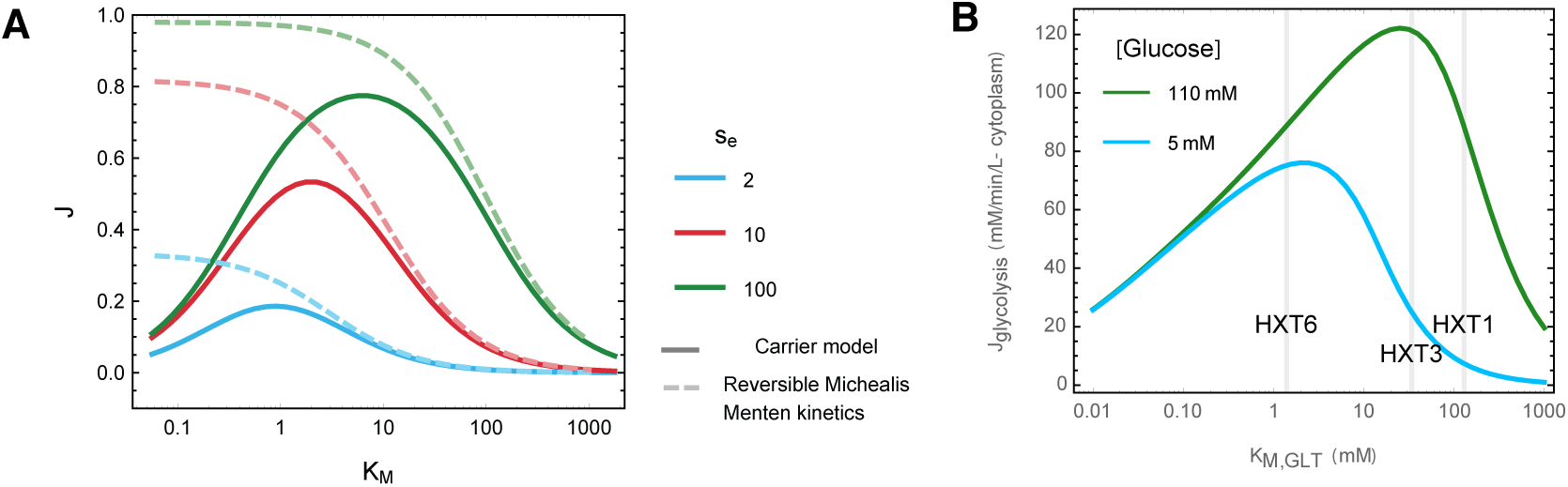
High affinities can reduce the net uptake rate in facilitated diffusion models A. Steady-state uptake rate *J* as a function of *K*_*M*_ of a facilitated diffusion model (solid lines) and with reversible Michaelis Menten kinetics (dashed lines), for different external substrate concentrations *s*_*e*_ and a constant internal sub-strate concentration *s*_*i*_ = 1. The *K*_*M*_ was varied by varying the substrate-transporter dissociation-constants *K*_*D*_, which for both rate equations affects the *K*_*M*_ but not the *k*_*cat*_. In a facilitated diffusion model very high affinities reduce the steady state flux, whereas for reversible Michaelis-Menten kinetics *J* monotonically decreases with *K*_*M*_. **B** Steady-state flux *J*_*glycolysis*_, as a function of the Michaelis-Menten constant of the glucose transporter *K*_*M,GLT*_, of a kinetic model of of *S. cerevisiae* glycolysis. At an extracellular glucose concentration of 110 mM (solid, black line) the low affinity HXT3 transporter (*K*_*M*_ = 34 mM) is roughly optimal, with *J*_*glycolysis*_ = 121 mM/ min /L cytosol, whereas the high affinity HXT6 transporter (*K*_*M*_ = 1.4 mM) attains *J*_*glycolysis*_ = 88.8 mM/ min /L cytosol. At low glucose concentrations (dashed, grey line) the high affinity transporter performs better.

The positive effects of reducing transporter affinity are only significant under certain conditions. For instance, if the intracellular substrate concentration is very low, substrate efflux is also low and thus hardly a problem. The intracellular substrate concentration depends on the kinetics of its uptake and consumption in metabolism in complex, non-intuitive ways. We therefore used a detailed kinetic model of *S. cerevisiae* glycolysis ([16] adapted from [17]) to test if under realistic conditions reducing the affinity significantly increases the net uptake rate. We calculated the steady-state glycolytic flux as a function of the *K*_*M*_ of the glucose transporter, *K*_*M,GLT*_ (Figure 3B). Indeed, at a high extracellular glucose concentration of 110 mM the low affinity HXT3 carrier (*K*_*M,GLT*_ ≈ 34 mM) attains a 36% higher glycolytic flux than the high affinity HXT6 carrier (*K*_*M,GLT*_ ≈ 1.5 mM). On the other hand, at low extracellular glucose concentration of 5 mM, the HXT6 transporter is expected to be (nearly) optimal. Note that we did not change the transporters *V*_*max*_, such that these difference arise purely from the difference in affinity.

### 2.4 Experimental observations do not discard any of the two hypotheses

Both the trade-off and the reduced efflux hypothesis can be tested using several single transporter strains with (wild-type or mutated) transporters differing in their *K*_*M*_. The absence of transporters with a high *K*_*M*_ and a high *k*_*cat*_ would support the trade-off hypothesis. Moreover, if this hypothesis is true, we would expect the wild-type transporters to be close to the Pareto front. (If a point lays on the Pareto front, there are no other points that have both a higher *k*_*cat*_ and a higher affinity.) A higher steady-state uptake rate at saturating substrate concentrations of cells with transporters with the same *k*_*cat*_ but a higher *K*_*M*_s would support the reduced efflux hypothesis.

Several labs have generated single glucose transporter strains in yeast. They expressed transporter constructs under constitutive promoters in a yeast glucose-transporter null-mutant. Elbing et al. generated chimera of the low-affinity HXT1 and the high-affinity HXT7 *S. cerevisiae* glucose carriers [10]. Kasahara et al. studied HXT7 [18, 19] and the low affinity HXT2 [20, 21, 22, 23] glucose carrier by either mutating specific residues or by constructing chimeras where specific trans-membrane segments from other carriers are used. For all these strains, the *K*_*M*_s and *V*_*max*_ have been measured. However, *k*_*cat*_ data are not available because quantitative measurements of carrier levels are technically demanding. Figure 4 shows the *K*_*M*_ and *V*_*max*_ measurements that we collected. The transporters from both labs show decreasing *V*_*max*_ with increasing affinity, and there are no constructs with both a low *K*_*M*_ and a high *V*_*max*_. In other words, there appears to be a *V*_*max*_-1/*K*_*M*_ Pareto-front (dashed gray line). Moreover, the wild-type transporters appear to sit on this Pareto front. Provided the expression levels of the carriers are similar, the *V*_*max*_ is a good reflection of the *k*_*cat*_, and these data support a rate-affinity trade-off. However, as we will discuss below, this assumption may not be valid.

**Figure 4:**
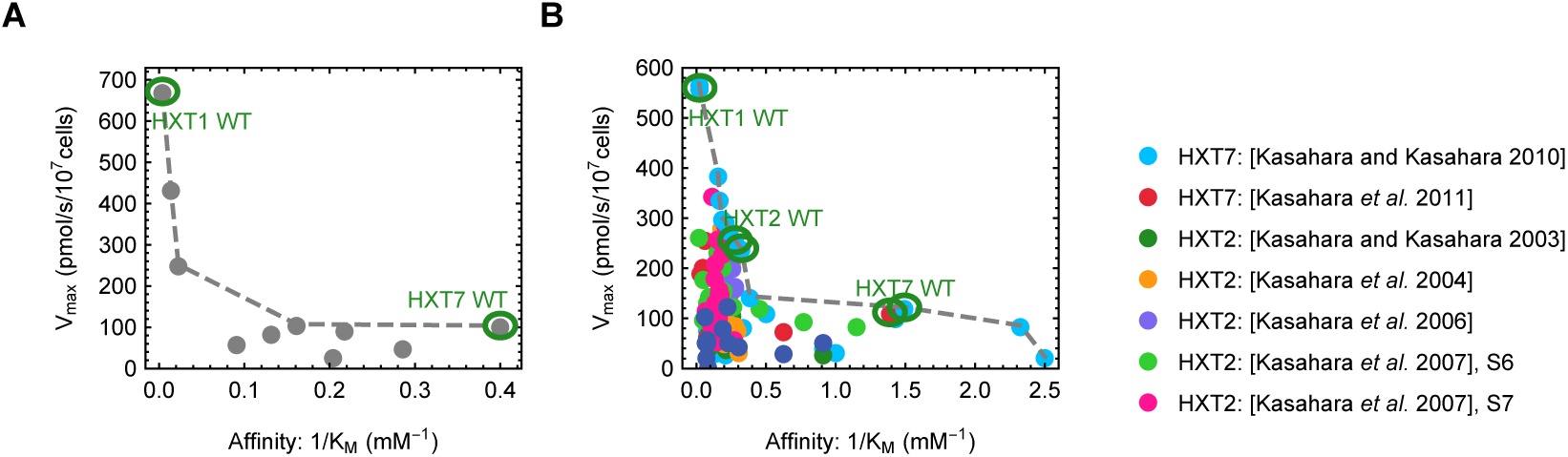
Experimental data suggest rate-affinity trade-off. Both panels show experimentally established affinities and rates of HXT mutants. **A** HXT1-HXT7 chimeras. Data taken from Elbing et al. [10]. (Converted from per mg of protein to per 10^7^ cells using 5 pg protein/cell, BioNumber ID:110550) **B** Different HXT2 and HXT7 mutants and chimeras constructed and characterised in the Kasahara lab. References are in the figure legend. The data suggest the existence of a *k*_*cat*_ 1/*K*_*M*_ Pareto-front (gray dashed line), provided the expression levels of all constructs is comparable. The wild-type transporters are located on this front.

To test our reduced-efflux hypothesis, we selected two constructs from the Kasahara lab that had a similar *V*_*max*_ but strongly differed in their *K*_*M*_: T213N (low affinity; *K*_*M*_ = 34 mM) and T213T (the wildtype HXT7 transporter with a high affinity; *K*_*M*_ = 0.72 mM) [19]. However, where those authors used transient expression, we integrated the gene into the genome behind the TDH3 promoter to obtain similar expression levels.

To evaluate our strains functionally we determined *V*_*max*_ and *K*_*M*_ values of the transporter with a 5s zero-trans influx uptake assay using radiolabelled glucose [24]. We confirmed the expected *K*_*M*_-values (see Table 1), but there was a 4-fold difference in *V*_*max*_ values between the two strains (Figure 5), considerably bigger than the 1.6-fold difference reported in Kasahara [19].

**Figure 5:**
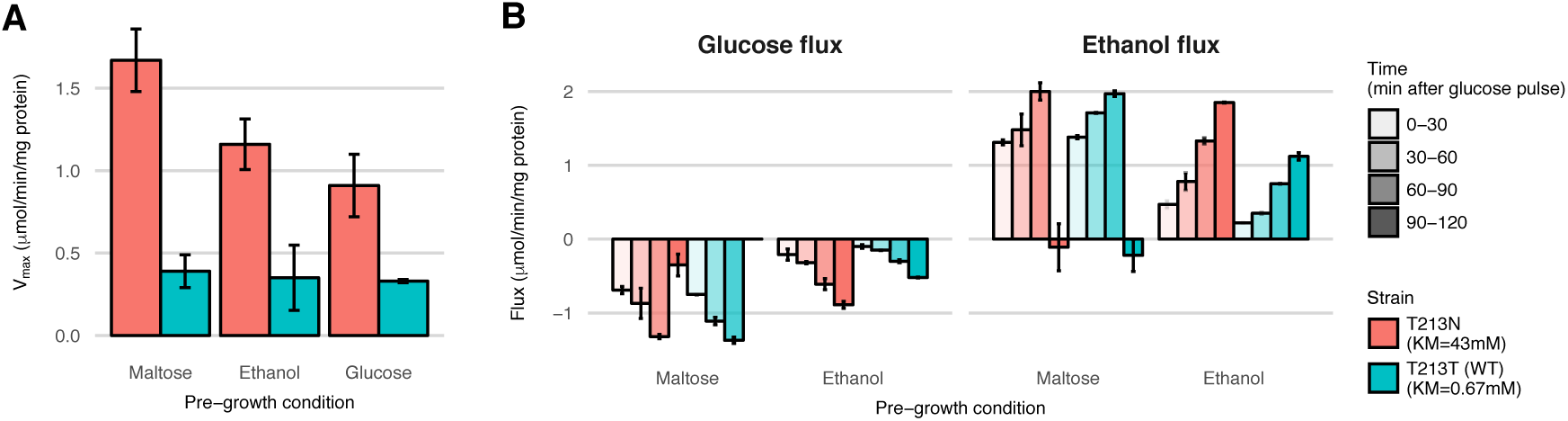
*V*_*max*_ values and steady-state fluxes for different pre-growth conditions. **A** *V*_*max*_ values were measured in a 5s zero-trans uptake assay with 200 mM glucose as substrate. Pre-growth was done on 2% of the carbon source indicated. Prior to the experiment cells were washed in medium without carbon source and incubated for 3' at 30 °C without a carbon source. Experiment was done in triplicate and shown are the averages with standard deviation. **B** Glucose and ethanol concentrations were measured during two hours at 30 °C following a glucose pulse. Protein concentration remained constant throughout the experiment. From this data, specific exchange fluxes were calculated. Shown is the average and standard deviation of two incubations per pre-growth condition.

**Table 1:**
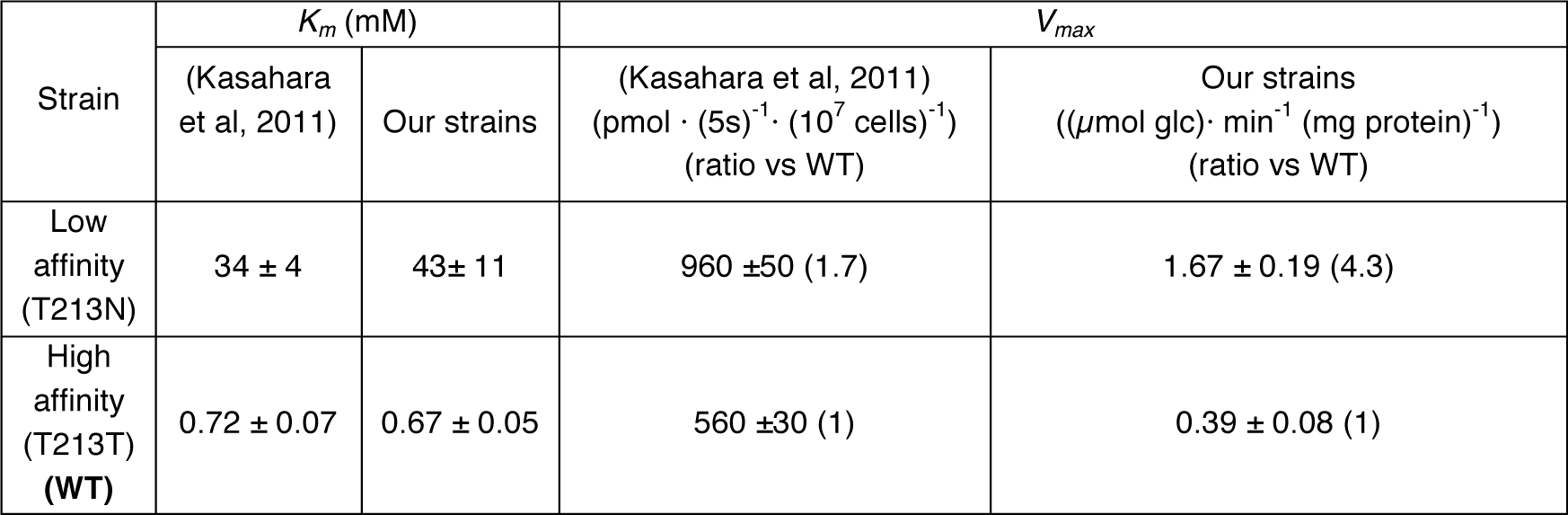
*K*_*M*_ and *V*_*max*_ values for glucose transport after pre-growth on maltose

| Strain | $K_m$ (mM) | | $V_{max}$ | |
| --- | --- | --- | --- | --- |
| | (Kasahara et al, 2011) | Our strains | (Kasahara et al, 2011)<br>($\text{pmol} \cdot (5\text{s})^{-1} \cdot (10^7 \text{ cells})^{-1}$)<br>(ratio vs WT) | Our strains<br>($(\mu\text{mol glc}) \cdot \text{min}^{-1} (\text{mg protein})^{-1}$)<br>(ratio vs WT) |
| Low affinity<br>(T213N) | $34 \pm 4$ | $43 \pm 11$ | $960 \pm 50$ (1.7) | $1.67 \pm 0.19$ (4.3) |
| High affinity<br>(T213T)<br>(WT) | $0.72 \pm 0.07$ | $0.67 \pm 0.05$ | $560 \pm 30$ (1) | $0.39 \pm 0.08$ (1) |

We tested whether there was an effect of the pre-growth conditions. Similar to Kasahara, the measurements above were obtained after pre-growth on maltose. Although we had carbon-starved the cells prior to the uptake assay, it could be that residual internal glucose from the conversion of maltose to glucose during pre-growth inhibits glucose uptake in our experiments. Along the lines of our hypothesis, this would affect the high-affinity strain more than the low-affinity strain and could result in a lower apparent *V*_*max*_ in the high-affinity strain. To overcome this potential problem, we grew the cells on the non-glycolytic substrate ethanol and again measured the *V*_*max*_ of glucose transport. Again, the low-affinity strain had a 4-fold higher *V*_*max*_ than the high-affinity strain. Pre-growth on glucose gave a lower *V*_*max*_ for the low-affinity strain, but still there was a 3-fold difference between the low-affinity and the high-affinity strain.

Steady-state flux experiments after a glucose pulse revealed further dynamics of uptake and glycolytic flux (Figure 5B). In the 2 hours after a glucose pulse, the glycolytic flux increases in each 30-minute interval. For cells that were pre-grown on maltose, the steady-state flux in the first 30 minutes exceeded the *V*_*max*_ we measured in the first 5s after the cells were exposed to glucose. This suggests that within 30 minutes the transport capacity was increased. Apparently, despite the genome integration, protein expression of these transporters depends on pre-growth medium, is highly dynamic, and might well differ between the different strains. Therefore, the *V*_*max*_ is not a good estimate of the *k*_*cat*_, which makes it hard to test the reduced efflux hypothesis and leaves some uncertainties to the experimental support for the trade-off hypothesis.

## 3 Conclusion and discussion

In this study we examined an existing hypothesis regarding the benefit of using low-affinity transporters, and proposed a new one. It has been postulated that a trade-off between rate and affinity exists, but so far theoretical as well as experimental evidence for this idea were lacking. We showed that for a facilitated diffusion uptake model there are theoretical arguments and experimental results in favour of this notion. However, there is no conclusive prove for this trade-off. We have also provided a novel explanation for the existence of low-affinity carriers, namely that they can reduce substrate efflux and thus enhance the net uptake rate.

The two hypotheses discussed here are not mutually exclusive. A decreased affinity might well be beneficial because it both raises the catalytic efficiency and reduces the substrate efflux. In fact, for the case of glucose uptake in yeast, a combination of a trade-off and the reduction of substrate efflux is probably the more complete explanation. The reduced efflux in that case might be the dominant effect for the high to intermediate affinity modulation. However, since for *K*_*M*_s above roughly 30 mM substrate efflux is likely negligible in any case, even lower affinities might be explained by the trade-off.

To test the reduced efflux hypothesis, we integrated the HXT7 isoforms made by Kasahara *et al* [19] into the genome of a yeast strain lacking glucose transporters. However, the HXT7 isoforms in our experiments did not only differ in *K*_*M*_, but also in their *V*_*max*_-values more than expected, and thus turned out to be less suitable to test the reduced efflux hypothesis. Despite identical sequence except for one mutation, identical nutrient conditions during pre-growth, and expression from the same promoter (tdh3), there is differential dynamics of the amount of transporters in the membrane in these two strains. Indeed, glucose-induced expression of the tdh-genes has been observed before [25, 26], and HXT7 is known to be degraded quickly in response to a glucose pulse [27]. Since it cannot be excluded that the transporter dynamics are affinity dependent (e.g. through the intracellular glucose levels), it is unclear if the *V*_*max*_ measurements presented in Figure 4 are a suitable proxy for *k*_*cat*_ values.

While there are observations that are in line with both hypotheses, the need for a solid experimental tests remains. The trade-off hypothesis could be verified experimentally with direct measurements of the *k*_*cat*_ values in addition to the *K*_*M*_ values. Those measurements would also allow for the selection of transporters for a renewed attempt to directly test the reduced efflux hypothesis. To tightly control transporter expression levels and other factors that influence the uptake capacity, measuring the uptake rate of transporters in artificial membranes might be an alternative, yet more artificial, test for the reduced efflux hypothesis.

Although direct experimental evidence turned out to be compromised by unresolved dynamics in glucose transporter activities, several observations support the reduced efflux hypothesis. First, in cells with high affinity transporters the steady-state glucose uptake rate at saturating extracellular glucose concentrations was up to 50% below the *V*_*max*_, and the measured intracellular glucose concentration was comparable to the transporter’s *K*_*M*_ [14]. For low affinity transporter, the steady-state flux matched the *V*_*max*_ and the intracellular glucose concentration was well below the *K*_*M*_ [14]. This indicates that intracellular glucose strongly inhibits uptake of the high but not of the low affinity transporter. Second, significant HXT-mediated glucose efflux has been observed in *S. cerevisiae* grown on maltose (maltose is intracellularly metabolized into two glucose molecules) [28]. Third, the affinity, but not the maximal uptake capacity, of glucose transport is modulated during growth on glucose [24].

Our reduced efflux hypothesis could also explain other phenomena besides low affinity uptake systems. For instance, the high affinity MCT1 and low affinity MCT4 lactate transporters in human tissue act as both importers and exporters, depending on the conditions [6]. MCT1 is typically expressed in cell types that import lactate as a substrate for oxidative phosphorylation, such as heart cells and red skeletal muscle, whereas MCT4 is predominantly expressed in cells that export excess lactate as a waste product of glycolysis, such as white skeletal muscle cells during heavy exercise [5]. Since the intracellular lactate concentration during heavy exercise is high, the low affinity MCF4 transporter may be favored because it wouldn’t be inhibited by low extracellular lactate.

To conclude, the idea that a lower affinity can result in a higher flux even at constant *k*_*cat*_ is perhaps counterintuitive, but is a direct consequence of the nature of transport processes, i.e. the absence of direct competition between extracellular and intracellular substrates. This enhanced product sensitivity is easily overlooked, and we expect it to be an important driver in systems where nutrient uptake is a relevant component of fitness.

## 4 Materials and Methods

**Strains and media** The *S. cerevisiae* strains used in this study are listed in Table S1. Under non-selective conditions, yeast was grown in complex medium (YPD) containing 10 g/L yeast extract, 20 g/L peptone and 20 g/L glucose. Selective media consisted of synthetic media (SM) containing 3 g/L KH_2_PO_4_, 0.5 g/L MgSO_4_ 7 H_2_O, 5 g/L (NH_4_)_2_SO_4_, 1 mL/L of a trace element solution and 1 mL/L of a vitamin solution [29]. Either maltose (SMM) or glucose (SMG) was used as sole carbon source (20 g/L). When necessary, the medium was supplemented to fulfil the auxotrophic requirements of the yeast strains, as previously described [30]. For the uptake experiments, strains were grown at 30 ◦C using defined mineral medium (CBS) [29] supplemented with required amino acids and 2% maltose or 2% ethanol as indicated.

**Strains and plasmids construction** EBY.VW4000, devoid of hexose transporters, was the platform strain for the constitutive expression of the HXT7 isoforms. Repair of the three auxotrophies of this strain (for tryptophan, leucine and histidine), construction of integrative vectors for the HXT7 isoforms and finally integration in EBY.VW4000 genome were the genetic modifications required.

*Repair of auxotrophies:* The three auxotrophies were repaired in a single transformation event (Figure S3). The TRP1 and HIS3 genes were PCR-amplified from the genomic DNA of *S. cerevisiae* CEN.PK113-7D using primers 5890_TRP1 fw, 5891_TRP1 rv, 2335_HIS3 fw and 2336_HIS3 rv, while the LEU2 gene was amplified from the pRS405 plasmid using primers 5892_LEU2 fw and 7052_LEU2 rv (Table S2). The primers used for these amplifications were designed to flank the integration cassettes with sequences homologous to the adjacent cassettes (SHR, [31]) or to the integration site in order to facilitate simultaneous assembly and integration in the CAN1 locus of EBY.VW4000 via homologous recombination. After transformation of the three integration cassettes to EBY.VW4000, transformants were grown on plates with SMM supplemented with uracil and subsequently replica-plated on SMM supplemented with uracil and L-canavanine (15 mg/L, [32]). A single colony was isolated and named IMX746. Proper assembly of the genes in IMX746 was confirmed by diagnostic PCR using the primers 5894_diag CAN1 fw, 5895_diag TRP1 rv, 5896_diag TRP1 fw, 5897_diag HIS3 rv, 5898_diag HIS3 fw, 5899_diag LEU2 rv, 5900_diag LEU2 fw and 5901_diag CAN1 rv (Table S2).

*Construction of integrative vectors for the HXT7 isoforms:* To ensure strong and constitutive expression, the HXT7 isoforms were cloned behind the *S. cerevisiae* TDH1 promoter. To this end, the HXT7 expressing plasmids were constructed by in vitro assembly (Gibson assembly) of three parts: a plasmid backbone obtained by PCR amplification of pRS406 using primers 6290_pRS406 fw and 6291_pRS406 rv, the TDH3 promoter PCR-amplified from the genomic DNA of CEN.PK113-7D with primers 6292_TDH2p fw and 6293_TDH3p rv and one of the two HXT7 isoforms amplified from the original Hxt7mnx-pVT_T213N and Hxt7mnx-pVT_T213T plasmids constructed by the Kasahara lab [19] using primers 6294_HXT7 fw and 6295_HXT7 rv. All primers are listed in Table S2. The PCR products were purified and, when the template for the reaction was a vector, digested with DpnI prior to assembly. Assembly of these parts yielded the integrative plasmids pUDI091 and pUDI093 (Table S3). Confirmation of the assembly was done by diagnostic PCR (with primers 6034_diag TDH3p fw, 1852_diag HXT7 rv, 1693_diag URA3 fw, 2613_diag TDH3p rv, 3795_diag ADH1t fw and 3228_diag pRS406 rv; all listed in Table S3) and by restriction analysis with BstBI.

*Genomic integration of HXT7 isoforms:* The strain IMX746 is auxotrophic for uracil due to a large insertion in the URA locus [33]. The cassettes pUDI091 and pUDI093, which carry the URA3 gene next to the HXT7 isoforms, were digested with the restriction enzyme StuI, that has a unique recognition site in the URA3 gene. Each of the linearized vectors were then transformed to the strain IMX746. The linearized vector was integrated at the URA3 locus via single crossover, resulting in the integration of a full copy of the URA3 gene and of the HXT7 isoform. Transformants were grown on plates containing SMG. Single colonies were isolated and named IMI335, for the strain containing the HXT7 T213N isoform and IMI337 for the strain containing the HXT T213T isoform. The sequences of the two HXT7 isoforms were confirmed by Sanger sequencing (Baseclear BV).

**Molecular biology techniques** PCR amplification with Phusion Hot Start II high-fidelity polymerase (Thermo Scientific, Waltham, MA) was performed according to manufacturer’s instruction using HPLC purified custom-synthesized oligonucleotide primers (Sigma-Aldrich). Diagnostic PCR was done with DreamTaq (Thermo Scientific) and desalted primers (Sigma-Aldrich). DNA fragments obtained by PCR were loaded on gel containing 1% (wt/vol) agarose (Thermo Scientific) and 1x Tris-acetate EDTA buffer (Thermo Scientific) and, when necessary, purified using the Zymoclean kit (Zymo Research, Irvine, CA). Propagation of plasmids was performed in chemically competent *E. coli* DH5α according to manufacturer instructions (Z-competent transformation kit, Zymo Research). Restriction analysis were performed with FastDigest enzymes DpnI, StuI and conventional enzyme BstBI (Thermo Scientific). Plasmids were isolated from *E. coli* with GenElute Plasmid kit (Sigma-Aldrich). Genomic DNA from yeast was extracted with YeaStar genomic DNA kit (Zymo Research). Cultures for transformation were grown overnight in complex medium with maltose. DNA concentrations of the cassettes for transformation were measured in a NanoDrop 2000 spectrophotometer (wavelength 260 nm, Thermo Fisher Scientific) and when necessary, pooled prior to transformation maintaining equimolar concentrations. Transformation to yeast was performed with the LiAc/ssDNA method described by Gietz et al. [34].

**Uptake experiments** *Mid-exponential phase cells for steady-state flux experiments:* Cells of two glycerol stocks of 1ml were washed with 10ml CBS medium (depending on the experiment with 2% maltose or 2% ethanol). The cells were cultured overnight in 10ml medium CBS with 2% maltose or 2% ethanol on a shaker at 30 ◦C and 200rpm. The culture was diluted to OD 0.2 in two separate 250 ml Erlenmeyer flasks, each with 50ml CBS with 2% maltose or 2% ethanol. Cultures were kept separate as duplicates. Cells were grown at 30 ◦C 200rpm until OD 4 and then diluted to OD 0.35 in 500ml CBS with 2% maltose or 2% ethanol in 1L erlenmeyer flasks. Cells were harvested at ~OD 4. Cells were pelleted at 1700g for 15 minutes and washed with 500ml CBS without carbon source. The pellet was re-suspended in CBS medium without carbon source to OD 40. The volume was measured and the cells were poured in a 100ml beaker for the experiment.

*Steady-state flux measurements:* The assay was performed at 30 ◦C. The cell suspension was stirred continuously. Samples for protein determination and metabolite concentration were taken prior to the start of the experiment. At time zero glucose was added to a final concentration of 4%. Samples were taken every 5 minutes during the first hour after addition of glucose and every 30 minutes during the second hour. For the glucose and ethanol concentration determination 0.1ml sample was added to 0.9ml 70% PCA and stored at -80 ◦C. After thawing, the pH of the samples was adjusted to ~4-5 with 185 µL 7M KOH. Samples were centrifuged and the supernatant was filtered through 0.22 µm syringe filter. The glucose and ethanol concentrations were determined with a Shimadzu LC20-AT HPLC and a Rezex (Phenomenex)™ column at 55 ◦C. The flow of the eluent (5mM H2SO4) was 0.5ml/min. Glucose and ethanol fluxes were calculated in µmol/(min mg-protein).

*Mid-exponential phase cells for zero trans influx uptake experiments:* Cells of a glycerol stock of 1ml were washed with 10ml medium CBS (depending on the experiment with 2% maltose or 2% ethanol). The cells were cultured overnight in 10ml medium CBS with 2% maltose or 2% ethanol on a shaker at 30 ◦C 200rpm. The culture was added to 50ml medium CBS with 2% maltose or 2% ethanol in a 250ml Erlenmeyer flasks and grown until ~OD 0.7, then the culture was diluted to ~OD 0.1 in three 250ml Erlenmeyer flasks, each with 50ml CBS with 2% maltose or 2% ethanol. Cultures were kept separate as triplicates. Cells were grown at 30 ◦C 200rpm until ~OD 3. Cells were pelleted at 1700g for 15 minutes and washed with 50ml CBS without carbon source. The pellet was re-suspended in CBS medium without carbon source to ~OD 80 and stored on ice. Prior to the uptake assay were put at 30 ◦C with aeration for 3 minutes to deplete intracellular glucose.

*Zero trans influx glucose uptake assay:* The zero-trans influx of ^14^C-labeled glucose was determined in a 5s assay, described by Walsh et al. [24] with the modifications of Rossell et al. [35]. The range of glucose concentrations used was between 0.25 and 200mM. Irreversible Michaelis-Menten equations were fitted to the data by non-linear regression.

**Protein samples and protein assay** For the steady-state flux experiments 0.25ml sample was added to 0.75ml CBS medium without carbon source and kept on ice. Cells were centrifuged for 2 minutes at maximal speed in a table centrifuge at 4 ◦C, washed with 1ml ice cold water, pelleted and resuspended in 1ml ice-cold 0.1M NaOH. For the zero-uptake experiments, 0.1ml of the cell suspension was added to a mix of 1ml NaOH 10M and 0.8ml demi water for the protein concentration. The samples were stored at -20 ◦C. After thawing the samples were boiled for 10 minutes at 110 ◦C, cooled down and centrifuged for 2 minutes at maximum speed. The supernatant was used for the protein determination with the Pierce BCA assay and compared to a dilution curve of BSA.

## Acknowledgements

This work was supported by NWO-VICI grant 865.14.005, and by projects funded by the Netherlands Genomics Initiative. We thank prof.Kasahara for kindly providing the HXT7 constructs, prof.Pascale Daran-Lapujade for help with integrating these into the genome, and Jose Gavaldá García for help with performing pilot experiments.

## Author contributions

EB, MTW and BT designed the research. EB and MTW developed the mathematical models and performed numerical simulations and theoretical analysis. PTC integrated the constructs into the genome. JRH and MJW performed the experiments. EB, MTW and JRH wrote the manuscript. All authors read and approved the manuscript.

## Footnotes

1 Here, we use Keq = 1 and assume symmetry between se and si release.

## Supplementary Text

### Symmetric transport model

#### Derivation of the rate equation of symmetric transport in terms of first order rate constants

The transporter can be in any of four states, the binding-site facing outward, with and without substrate bound, *es*_*e*_ and *e*_*e*_, respectively, and inward facing with and without substrate bound, *es*_*i*_ and *e*_*i*_. Assuming that binding and unbinding is much faster than the movement of the binding site over the membrane, we can use the quasi-steady state approximation for the fraction of carriers that have substrate bound to them, both inside and outside,

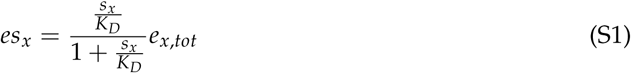

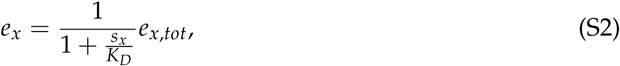

where *K*_*D*_ is the substrate-transporter dissociation constant and *e*_*x*,*tot*_ = *es*_*x*_ + *e*_*x*_ is the total number of transporters with their binding site facing the *x* site of the membrane (i.e. *x* = *e* or *x* = *i*).

By definition, in steady state *e*_*x,tot*_ is constant. This gives rise to the equality *k*_2_*es*_*e*_ + *k*_4_*e*_*e*_ = *k*_2_*es*_*i*_ + *k*_4_*e*_*i*_. Defining

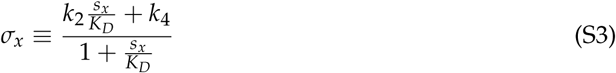

and solving the steady state condition gives an expression for the total amount of outward and inward facing carriers, normalized to the total amount of transporters:

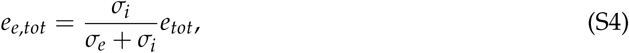

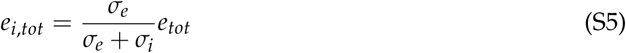

The net uptake rate is than given by:

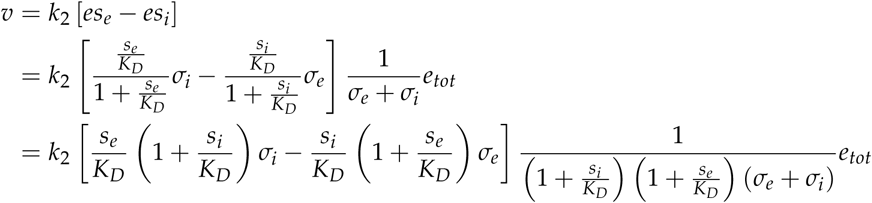

Filling in the σs, the term within the square brackets reduces to

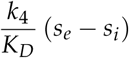

and the denominator to

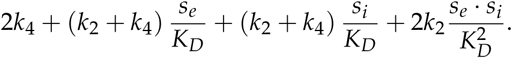

Hence, the rate equation in terms of the first order rate constants is given by

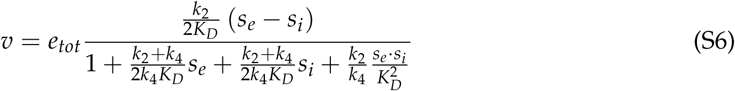

Defining the macroscopic kinetic parameters

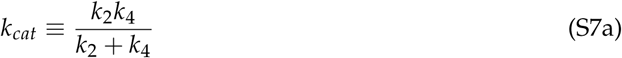

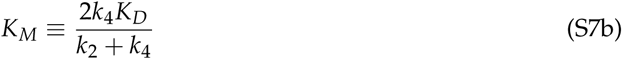

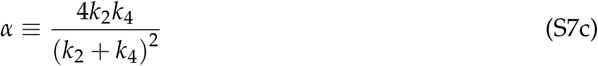

than gives the rate equation

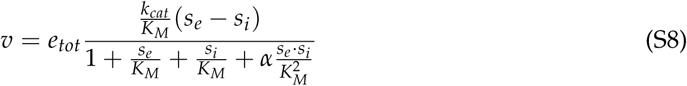

#### Derivation of the optimal affinity,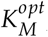

In order to find the optimal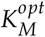 we simply take the derivative of v with respect to *K*_*M*_ and set that to zero. Since we have

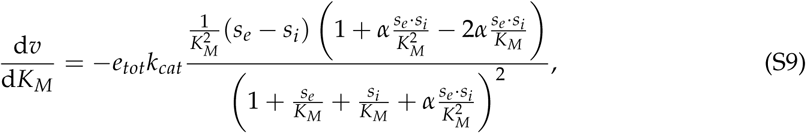

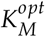 is found by solving

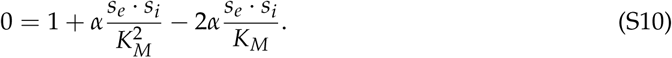

The physical (i.e. positive) solution to this quadric equation is

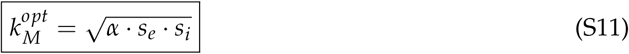

In comparison, the reversible Michaelis-Menten rate equation^1^,

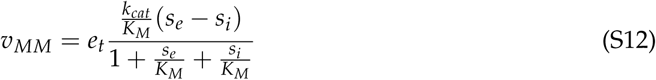

does not have an optimal affinity. Reducing the *K*_*M*_ will always increase the rate.

### The non-symmetric carrier model

In this section we study if the conclusions in the main text hold when the assumptions underlying the simplification of the rate equation are dropped. The more general rate equation describing the net transport rate of a facilitated diffusion process takes the following form (equation IX-46 in [36]):

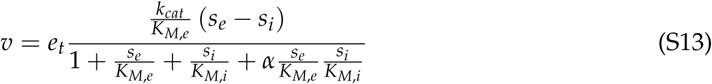

where the macroscopic kinetic parameters, *k*_*cat*_, *K*_*M,e*_, *K*_*M,i*_ and α, can be expressed in terms of the first order rate constants.

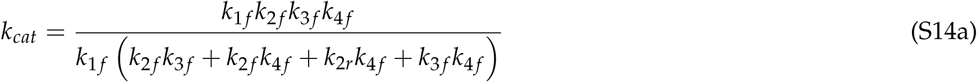

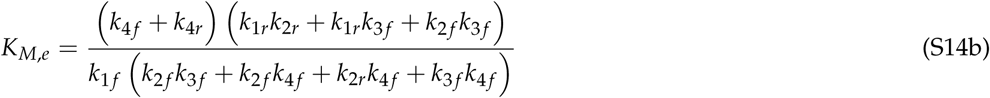

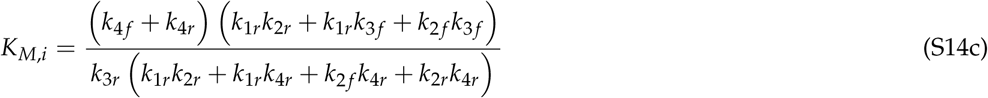

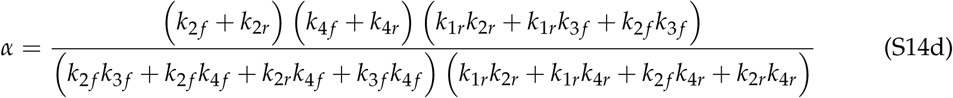

Furthermore, the first order rate equations are related through the equilibrium constant. Since we are considering facilitated diffusion, no free energy dissipation is coupled to the transport process,*i.e. K*_*eq*_ = 1, and we have:

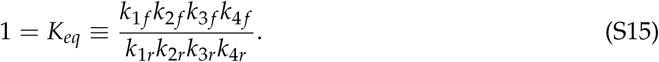

This poses a constraint on the first order rate constants. Practically, this means that a mutation that affects *e.g.* the strength of extracellular substrate to carrier binding must also affect some of the other steps in the transport cycle (*e.g.* intracellular substrate release). Combined with the complicated dependency of the macroscopic parameters on the first order rate constants, an analytical approach is unfeasible. We therefore employed a parameter sampling approach to gauge to what extent our conclusions about the rate-affinity trade-off and the substrate efflux hypothesis are valid for this more general rate equation.

#### The rate-affinity trade-off

As in the main text, the parameter sampling approach gives a mixed picture about the theoretical underpinnings of the rate-affinity trade-off. Figure S1 shows scatterplots of *k*_*cat*_ versus affinity (defined as 1/*K*_*M,e*_) for randomly sampled sets of first order rate constants, with a number of different constraints assumed for some of these. In the absence of any constraints, there is a clear negative correlation between the *k*_*cat*_ and the affinity (figure S1A). However, this is not a true Pareto-front, as it appears as though there is always a possibility that the *k*_*cat*_ is enhanced without reducing the affinity (or vice verse). The fact that not the whole *k*_*cat*_ 1/*K*_*M*_ space is filled is due to the finite numbers of samples rather than due to a true constraint. On the other hand, if we assume that there is a (biophysical) limitation on the rate of substrate-transporter binding (*k*_1 *f*_), (*e.g.* the diffusion limit), we do find a true Pareto front (figure S1B), the location of which depends on the actual maximal *k*_1 *f*_ -value. However, this conclusion does not hold if other rate constants are assumed to have some biophysical limit, as shown by the examples of restricted *k*_2 *f*_ (figure S1C) or a restricted substrate-transporter dissociation constant *K*_*D,e*_ (*≡ k*_1*r*_/*k*_1 *f*_, figure S1D). All in all, there are reasonable theoretical arguments to be made for a rate-affinity trade-off, but the logic is not water-tight.

#### Enhanced uptake by reduced efflux

In the analysis in the main text above we made the biologically motivated, simplifying assumption that the transporter is symmetric. However, our reasoning does not critically dependent on this symmetry, since it is a general property of this scheme that the substrate and product bind to different states of the transporter. Moreover, since all first order rate are interdependent through constraint (S15), *K*_*M,i*_ and *K*_*M,e*_ are expected to be correlated. To test this, we randomly sampled all first order rate constants from a log-normal distribution and rescaled them such that *K*_*eq*_ = 1 and *k*_*cat*_ = 1 (for details, cf. Appendix 4). These parameters were used to calculate the *K*_*M,e*_, *K*_*M,i*_ and the net steady state uptake rate *J* under conditions of high an low external substrate (*s*_*e*_ = 100 and *s*_*e*_ = 1, respectively). The results are depicted in figure S2. Indeed, *K*_*M,e*_ and *K*_*M,i*_ are correlated, albeit not strongly (Spearman correlation = 0.64, figure S2A). More importantly, however, there appears to be an *s*_*e*_-dependent optimal affinity (figure S2B). Furthermore, the set of parameters that has the highest *J* under low substrate conditions, performs relatively poorly under high substrate conditions (large, light red dot), and vice verse (blue dot).

### Parameter sampling procedure

Sets of parameters were constructed by drawing the first order rate constant randomly from a log-normal distributions. This was done in a way such that constraint (S15) is satisfied. These parameter sets were used to calculate the steady state uptake rate (given by equation (S13)) and macroscopic kinetic parameters (given by equation (S14)). The parameter sampling was performed in Wolfram Mathematica 9.0 using the functions RandomVariate and LogNormalDistribution, which has the probability density function (PDF):

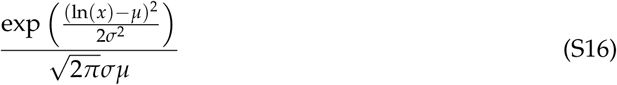

#### Rate affinity trade-off

To generate the data depicted in Figure S1, for each subfigure 10 000 parameter sets were constructed. Each parameter set was constructed in the following way:

- Two sets of four numbers, *χ* ≡ {*x*_1_, *x*_2_, *x*_3_, *x*_4_} and {*y*_1_, *y*_2_, *y*_3_, *y*_4_} were randomly drawn from a log-normal distribution given by the PDF (S16) with *µ* = 0 and *σ* = 2.
- For **figure S1A**, there are no restrictions on individual rate constants. The set of forward rate constants, *K*_*f*_ ≡ {*k*_1 *f*_, *k*_2 *f*_, *k*_3 *f*_, *k*_4 *f*_} is just given by the first set, *K* _*f*_ = *X*. To get the reverse rate constants *K*_*r*_ ≡ {*k*_1*r*_, *k*_2*r*_, *k*_3*r*_, *k*_4*r*_}, *Y* needs by be rescaled by a factor

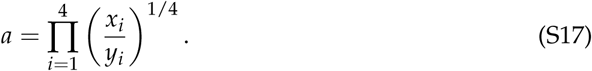

This is to ensure that *K*_*eq*_ = 1. Hence, *K*_*r*_ = a · *Y*.

- For **figure S1B** we also need to ensure that *k*_1 *f*_ is restricted to some constant value c. We set the first order rate forward constants to 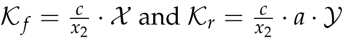 Similarly, for **figure S1C** we need to restrict *k*_2*f*_ to *c*. We use 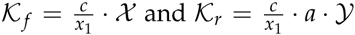
- For **figure S1D**, the *K*_*D,e*_ is fixed to a constant value *c*. Here, we used for the forward rate constant simply *K*_*f*_ = *X*. Since *K*_*D,e*_ ≡ *k*_1*r*_/*k*_1*f*_, we set *k*_1*r*_ = *c* · *x*_1_. Defining β = 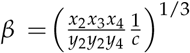 and setting {*k*_2*r*_, *k*_3*r*_, *k*_4*rg*_} = β · {*y*_2_, *y*_3_, *y*_4*g*_ additionally ensures *K*_*eq*_ = 1.

#### Enhanced uptake by non-symmetric low affinity transporters

Figures S2 A and B are constructed from the same 10000 parameter sets. Each set was constructed such that *k*_*cat*_ = 1 and *K*_*eq*_ = 1 as follows:

- Two sets of four numbers, *χ* ≡ {*x*_1_, *x*_2_, *x*_3_, *x*_4_} and *Y* ≡ {*y*_1_, *y*_2_, *y*_3_, *y*_4_} were randomly drawn from a log-normal distribution given by the by the PDF (S16) with *µ* = 0 and *σ* = 2.
- The set *Y’*, is obtained by rescaling *Y* by a factor

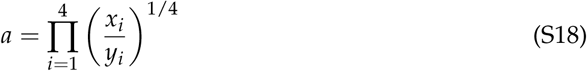

to obtain 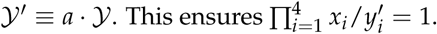
- Both sets are rescaled by a factor 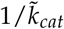, as defined in equation (S7a) with *k*_*if*_ → *x*_*i*_ and 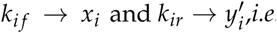

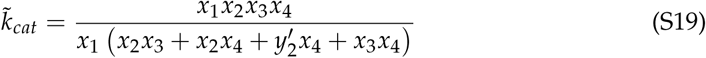 This rescaling is done to ensure that *k*_*cat*_ = 1 for each parameter set. The first order rate 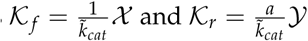

## Supplementary Tables

**Table S1:** Saccharomyces cerevisiae strains used in this study

| Name | Genotype | Relevant phenotype | Origin |
| --- | --- | --- | --- |
| CEN.PK113-7D | *MATa URA3 HIS3 LEU2 TRP1 MAL2-8c SUC2* | Prototrophic control strain | Euroscarf |
| EBY.VW4000 | *MATa ura3 his3 leu2 trp1 MAL2-8c SUC2 hxt13Δ::loxP hxt15Δ::loxP hxt16Δ::loxP hxt14Δ::loxP hxt12Δ::loxP hxt9Δ::loxP hxt11Δ::loxP hxt10Δ::loxP hxt8Δ::loxP hxt514Δ::loxP hxt2Δ::loxP hxt367Δ::loxP gal2Δ stl1v::loxP agt1Δ::loxP ydl247wΔ::loxP yjr160cΔ::loxP* | HXT null strain, auxotrophic for uracil, leucine, histidine and tryptophan. Does not grow on glucose as sole carbon source | Wieczorke et al. (1999) |
| IMX746 | *MATa ura3 MAL2-8c SUC2 hxt13Δ::loxP hxt15Δ::loxP hxt16Δ::loxP hxt14Δ::loxP hxt12Δ::loxP hxt9Δ::loxP hxt11Δ::loxP hxt10Δ::loxP hxt8Δ::loxP hxt514Δ::loxP hxt2Δ::loxP hxt367Δ::loxP gal2Δ stl1v::loxP agt1Δ::loxP ydl247wΔ::loxP yjr160cΔ::loxP can1::LEU2-HIS3-TRP1* | HXT null strain with a single auxotrophy for uracil | This study |
| IMI335 | *MATa MAL2-8c SUC2 hxt13Δ::loxP hxt15Δ::loxP hxt16Δ::loxP hxt14Δ::loxP hxt12Δ::loxP hxt9Δ::loxP hxt11Δ::loxP hxt10Δ::loxP hxt8Δ::loxP hxt514Δ::loxP hxt2Δ::loxP hxt367Δ::loxP gal2Δ stl1v::loxP agt1Δ::loxP ydl247wΔ::loxP yjr160cΔ::loxP can1::LEU2- HIS3-TRP1 ura3::HXT7_T213N-URA3.* | Prototrophic HXT null strain expressing HXT7_T213N | This study |
| IMI337 | *MATa MAL2-8c SUC2 hxt13Δ::loxP hxt15Δ::loxP hxt16Δ::loxP hxt14Δ::loxP hxt12Δ::loxP hxt9Δ::loxP hxt11Δ::loxP hxt10Δ::loxP hxt8Δ::loxP hxt514Δ::loxP hxt2Δ::loxP hxt367Δ::loxP gal2Δ stl1v::loxP agt1Δ::loxP ydl247wΔ::loxP yjr160cΔ::loxP can1::LEU2-HIS3-TRP1 ura3::HXT7_T213T-URA3.* | Prototrophic HXT null strain expressing HXT7_T213T | This study |

**Table S2:**
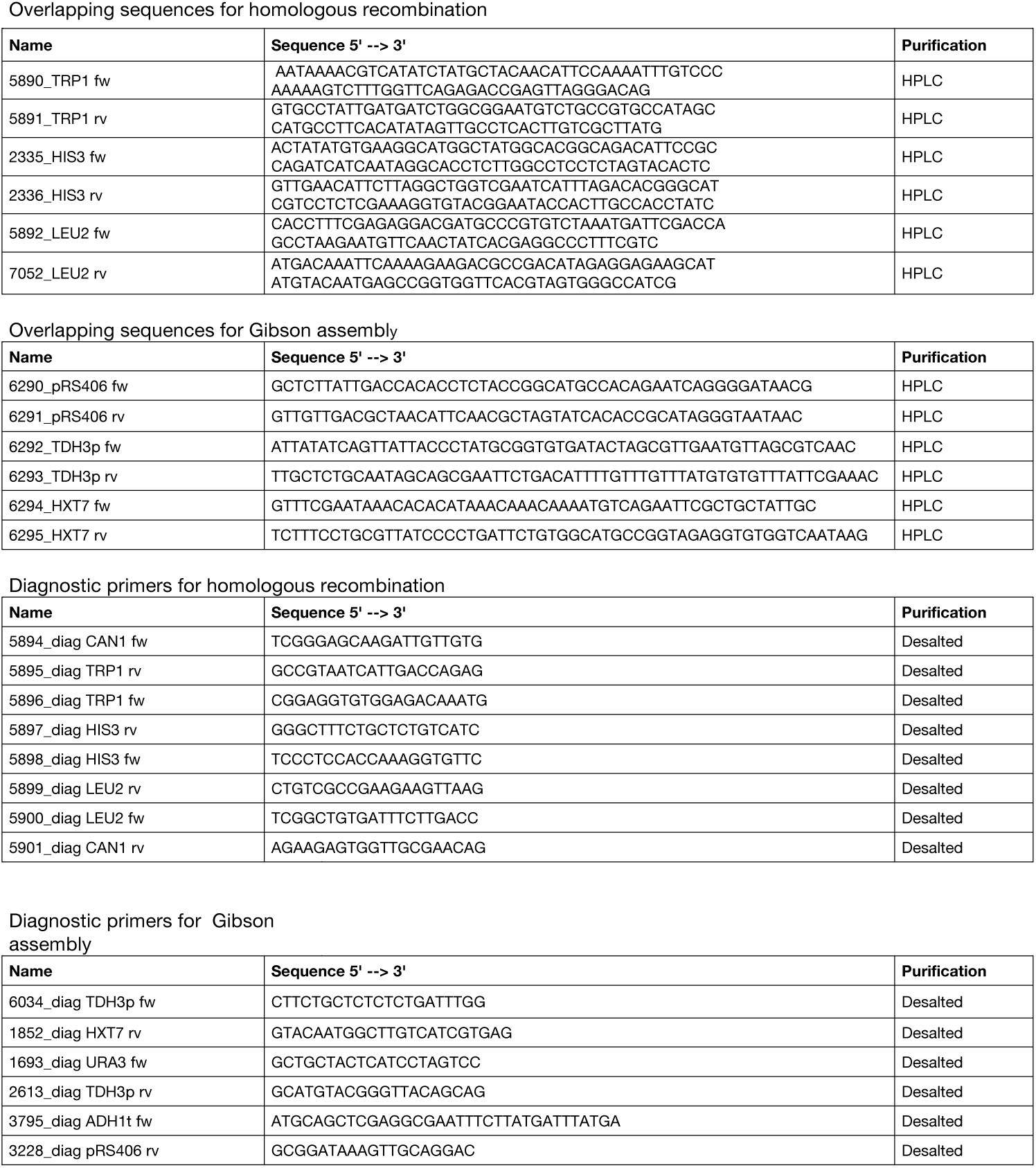
Primers used in this study

**Table S3:**
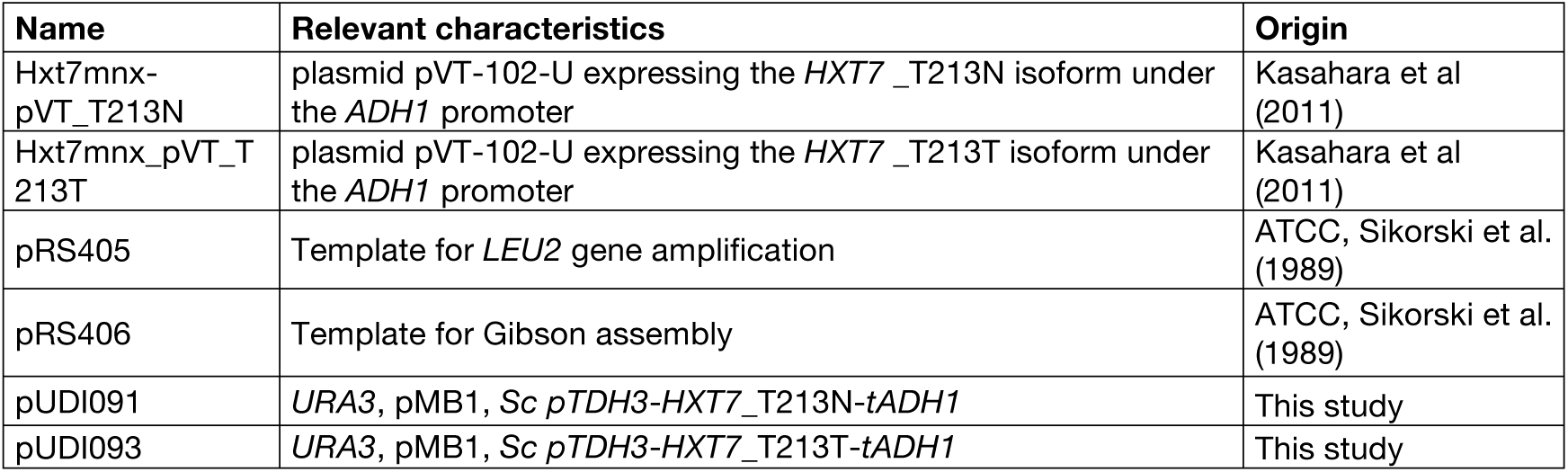
Plasmids used in this study

| **Name** | **Relevant characteristics** | **Origin** |
| --- | --- | --- |
| Hxt7mnx-pVT_T213N | plasmid pVT-102-U expressing the *HXT7*_T213N isoform under the *ADH1* promoter | Kasahara et al (2011) |
| Hxt7mnx_pVT_T213T | plasmid pVT-102-U expressing the *HXT7*_T213T isoform under the *ADH1* promoter | Kasahara et al (2011) |
| pRS405 | Template for *LEU2* gene amplification | ATCC, Sikorski et al. (1989) |
| pRS406 | Template for Gibson assembly | ATCC, Sikorski et al. (1989) |
| pUDI091 | *URA3*, pMB1, *Sc pTDH3-HXT7_T213N-tADH1* | This study |
| pUDI093 | *URA3*, pMB1, *Sc pTDH3-HXT7_T213T-tADH1* | This study |

**Figure S1:**
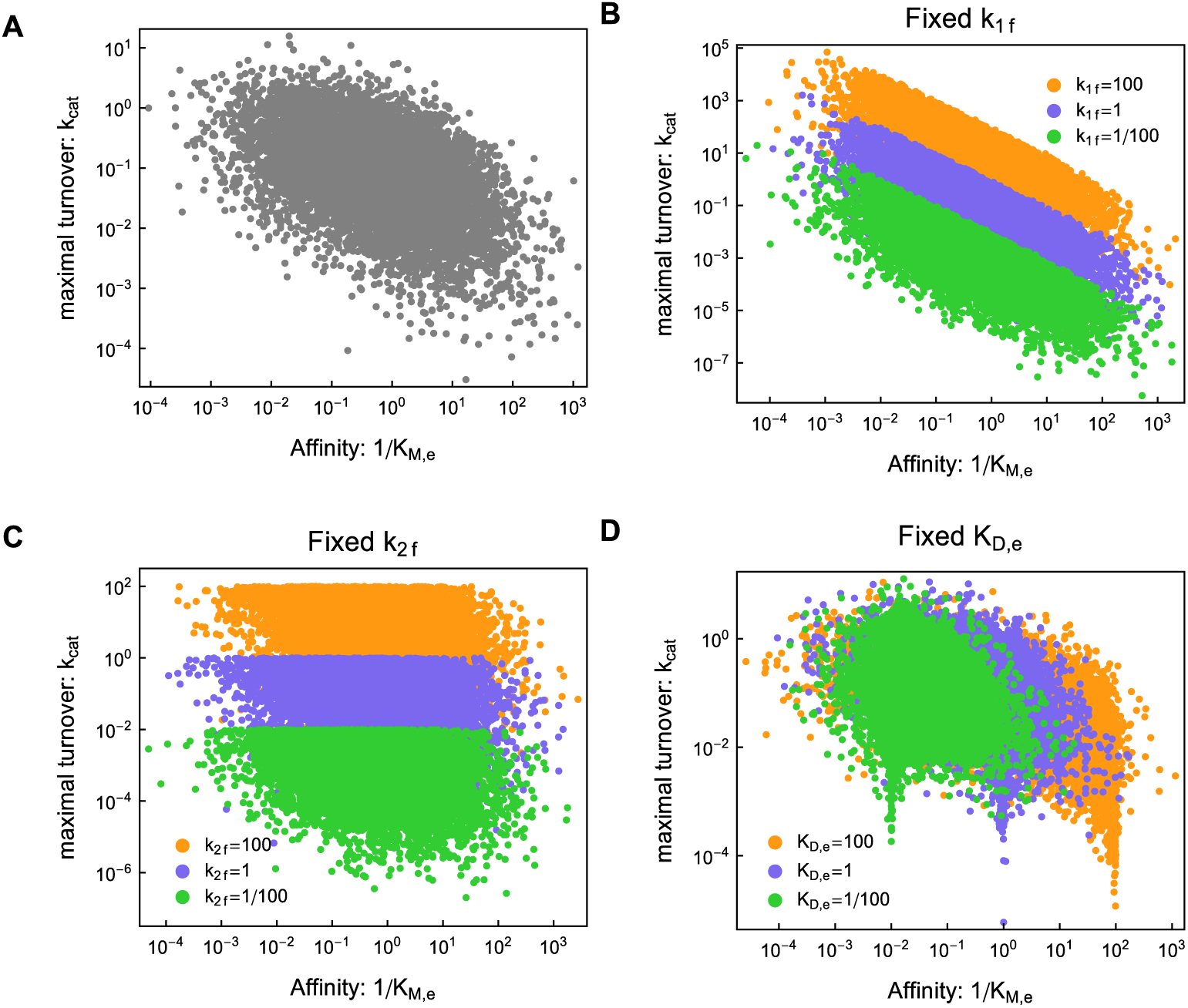
Parameter sampling gives inconclusive picture of potential rate-affinity tradeoff. See Supplementary Information for explanations and details on the parameter sampling procedure.

**Figure S2:**
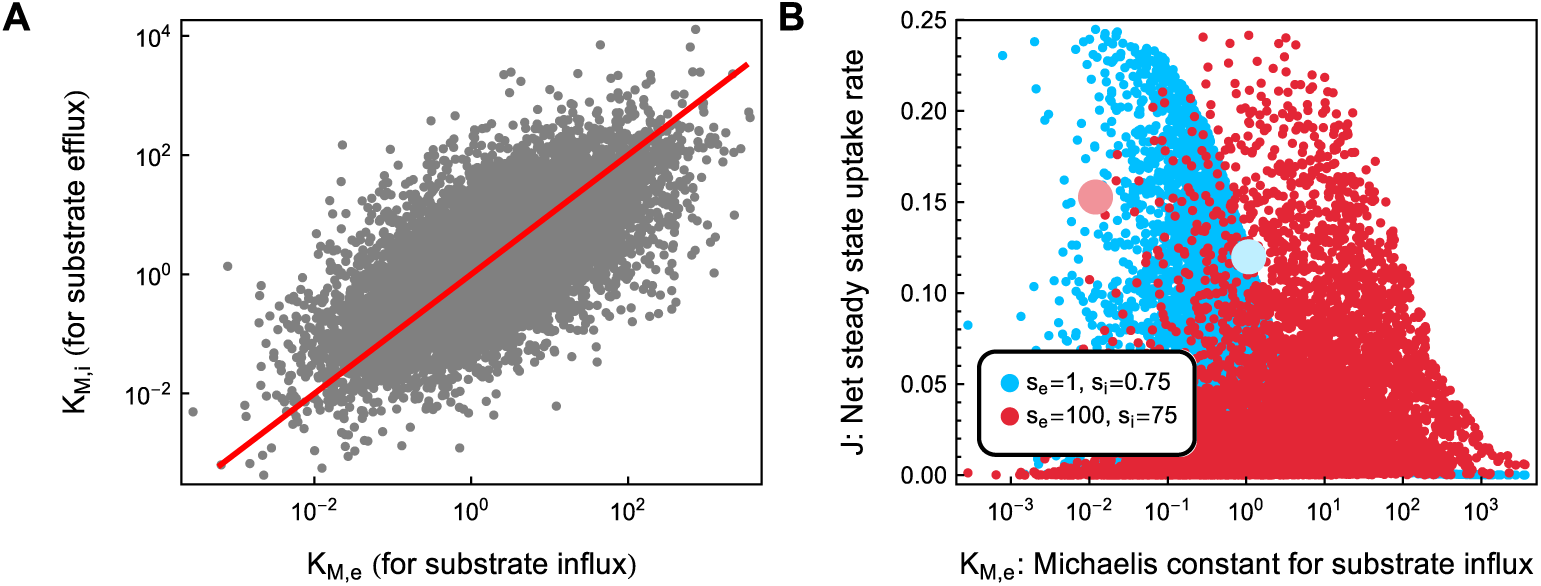
Parameter sampling confirms enhanced uptake by non-symmetric low affinity carriers. The large red (blue) dot in panel **B** indicate the set parameters that are optimal at low *s* (high *s*). See Supplementary Information for details on the parameter sampling procedure

**Figure S3:**
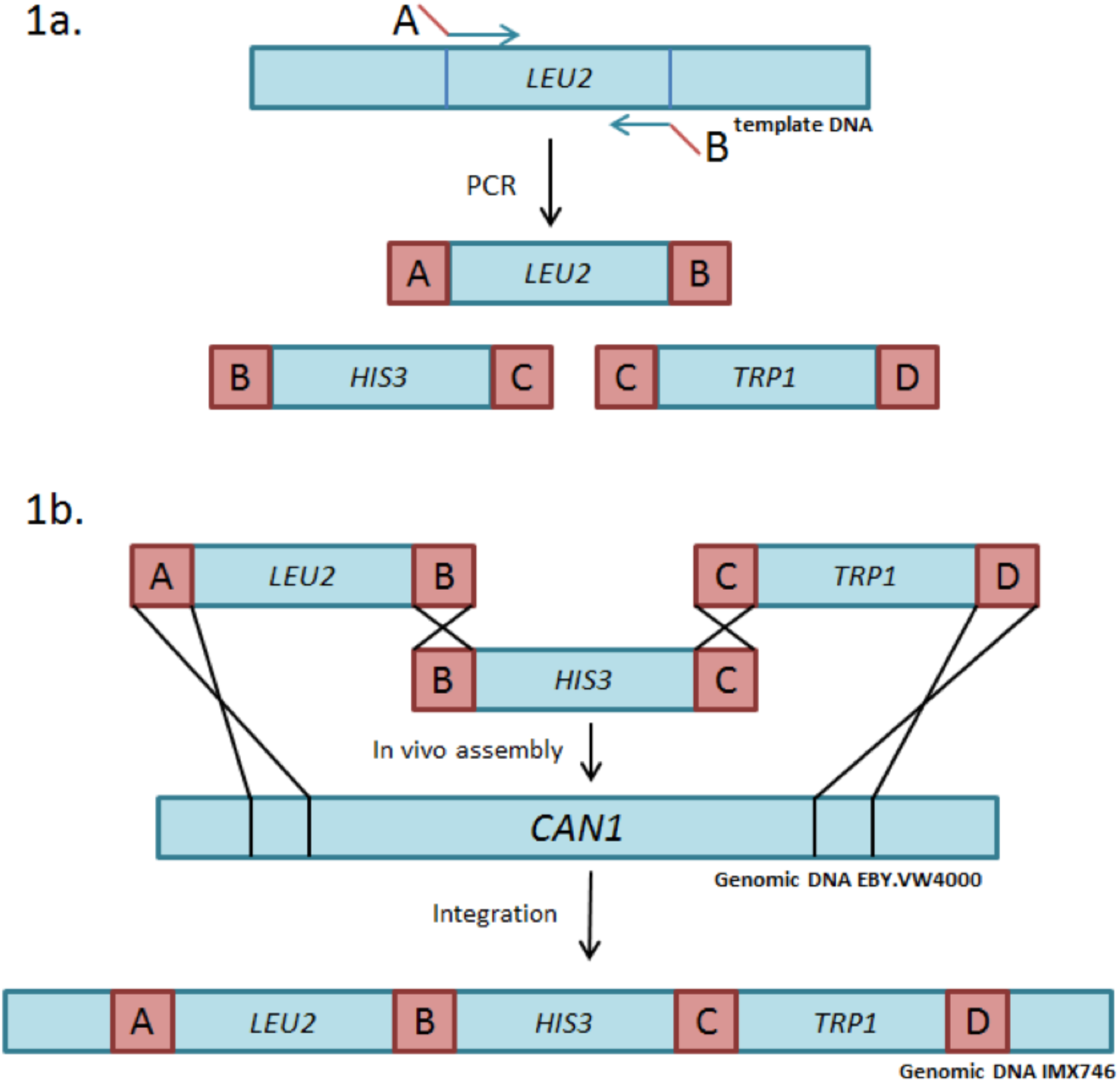
Repair of auxotrophies in the EBY.VW4000 strain. **A** PCR amplification of the LEU2, HIS3 and TRP1 genes with sequences overlapping either the adjacent cassettes or the integration site. **B** In vivo assembly of the marker cassettes and integration in the CAN1 locus of EBY.VW4000.

